# Unique habitat and macroinvertebrate assemblage structures in spring-fed stream: a comparison among clastic lowland tributaries and mainstreams in northern Japan

**DOI:** 10.1101/2020.05.17.101014

**Authors:** Masaru Sakai, Katsuya Iwabuchi, David Bauman

## Abstract

The stable flow and temperature regimes of spring-fed streams are distinct from the dynamic regimes of other streams. We investigated differences in habitat and macroinvertebrate assemblages among three stream types (spring-fed tributary, non-spring-fed tributary and mainstream) in a clastic lowland of northern Japan. Current velocity was the slowest in the spring-fed reach, where the percent of fine sediment deposition was also 3.8–11.4 times higher than in the other stream types. The standing stock of detritus was also greater in the spring-fed reach. These results suggest that the stable flow regime in the spring-fed stream leads to the accumulation of fine sediment and detritus on the streambed. Oligochaeta and chironomids, which are burrower-gatherers, were remarkably abundant in the spring-fed reach. Total macroinvertebrate abundance was 3.8–12.2 times greater in the spring-fed reach than in the other stream types. Sprawler-grazer ephemeropterans were the most abundant in the mainstream reaches, likely due to higher primary productivity. *Allomyia* sp, which depend on cool spring-fed habitats, were found only in the spring-fed reach. The indicator species analysis also indicated multiple taxa of detritivores and *Allomyia* sp. for the spring-fed tributary. The macroinvertebrate assemblage in the spring-fed reach was characterized by numerous burrowers, collector-gatherers, and crenobiont taxa, highlighting the uniqueness and its contribution to enhance beta diversity in river networks.

## Introduction

Groundwater flow into river ecosystems depends on underground flow pathways and aboveground topographic features (Allan and Castillo 2007). Although apportioning stream water to runoff and groundwater sources is generally difficult, streams that are sustained by a significant amount of permanent groundwater discharge from a spring mouth are typically referred to as spring-fed streams (Cantonati et al. 2010). Because groundwater flow is rarely affected by rapid fluctuations resulting from rainfall, snowmelt, or seasonal and diurnal changes in air temperature, spring-fed streams are characterized by stable water flow and temperature regimes (Mattson et al. 1995; Sear et al. 1999; Lusardi et al. 2016). Thus, spring-fed streams are distinct from the dynamic flow and temperature regimes that characterize non-spring-fed streams.

Flow regime is one of the most influential factors determining the structure of streambed substrates (Davis and Barmuta 1989). The frequency and magnitude of high flows are directly related to sediment runoff; stream substrate components are sequentially flushed from smaller to larger particles under high flows (Richards 2004). Streams with frequent high flows typically have a coarse streambed structure relative to those with less variable flow regimes. The substrates of spring-fed streams are dominated by fine sediments, owing to their stable flow regimes (Sear et al. 1999). Spring-fed streams in lowland regions may have particularly fine streambeds due to low channel gradients. Streambed particle size is a major determinant of the benthic macroinvertebrate community and its structure (Richards and Bacon 1994; Wood and Armitage 1997). For example, the relative density of burrowers and gatherers may increase with fine sediment deposition (Rabení et al. 2005). These groups are well suited to depositional habitats because these habitats provide organic matter as well as burrowing spaces. Lowland spring-fed streams may therefore house unique community assemblages, dominated by burrowers and gatherers, relative to adjacent non-spring-fed streams that have less fine sediment content.

Water temperature in spring-fed streams is highly stable; it is typically cold in summer and warm in winter (Mattson et al. 1995). Previous studies have reported that these stable water temperature regimes provide refugia for aquatic fauna to avoid severe hot (Nakagawa et al. 2015) and cold seasonal conditions (Inoue and Ishigaki 1968). Moreover, Sun et al. (2020) suggested that most indicator species of spring-fed habitats may be stenothermal. This further suggests that the narrow range of water temperatures in spring-fed streams may lead to the formation of unique assemblages of crenobiont and/or crenophilous species (Lusardi et al. 2016).

The stable flow and temperature regimes of spring-fed streams provide unique habitat and macroinvertebrate assemblage structures and subsequently contribute to beta diversity within river networks. However, studies on spring ecosystems have advanced mainly in carbonate geologies (e.g., karst) with dense spring habitats (Mattson et al. 1995; Wood et al. 2005; von Fumetti et al. 2017; von Fumetti and Blattner 2017), and the ecosystem functions are poorly understood in other geologies. This is perhaps related to sparse distributions of springs and difficulties to prepare replicable sampling design in such geologies. Although more than 16,000 spring-fed sites have been recognized in Japan (Ministry of Environment 2020), functional aspects of spring-fed streams are still elusive in any geological conditions. We assessed differences in habitat characteristics and macroinvertebrate assemblages between spring-fed and adjacent non-spring-fed streams in a clastic lowland, one of the dominant geological conditions in Japan. Our aim was to investigate potential processes explaining these differences, detect potential indicator species of different stream types, and to contribute to the understandings of faunal and functional responses of macroinvertebrates to the unique habitats of spring-fed tributaries.

## Materials and methods

### Study area

We assessed streams in the Shubuto River system, located in Kuromatsunai, Hokkaido, Japan (42.64°N, 140.34°E). The Shubuto River basin comprises 367 km^2^ of montane and lowland areas; our study focused on the lowland region (Fig. 1). Water quality is suitable for most freshwater organisms: dissolved oxygen > 95% in degrees of saturation, biochemical oxygen demand is 0.2-1.7 mg/L and ammonia concentration < 0.05mg/L (Miyazaki and Terui 2016). The geology is dominated by sandstone and mudstone with Cainozoic fossil seashells and tuff. Mean annual precipitation and air temperature, obtained from the Kuromatsunai automated meteorological data acquisition system station (located 4 km northwest of the site) were 1615.8 mm and 7.5°C, respectively, between 2009 and 2018. Dominant tree species in the riparian zones of the study area were *Salix* spp. and *Quercus crispula*, and dominant understory plants were *Sasa kurilensis* and *Reynoutria sachalinensis*. High flows generally occur in non-spring-fed streams during summer to autumn rainfalls and snowmelt season in early spring (Fig. 2).

**Fig. 1.**
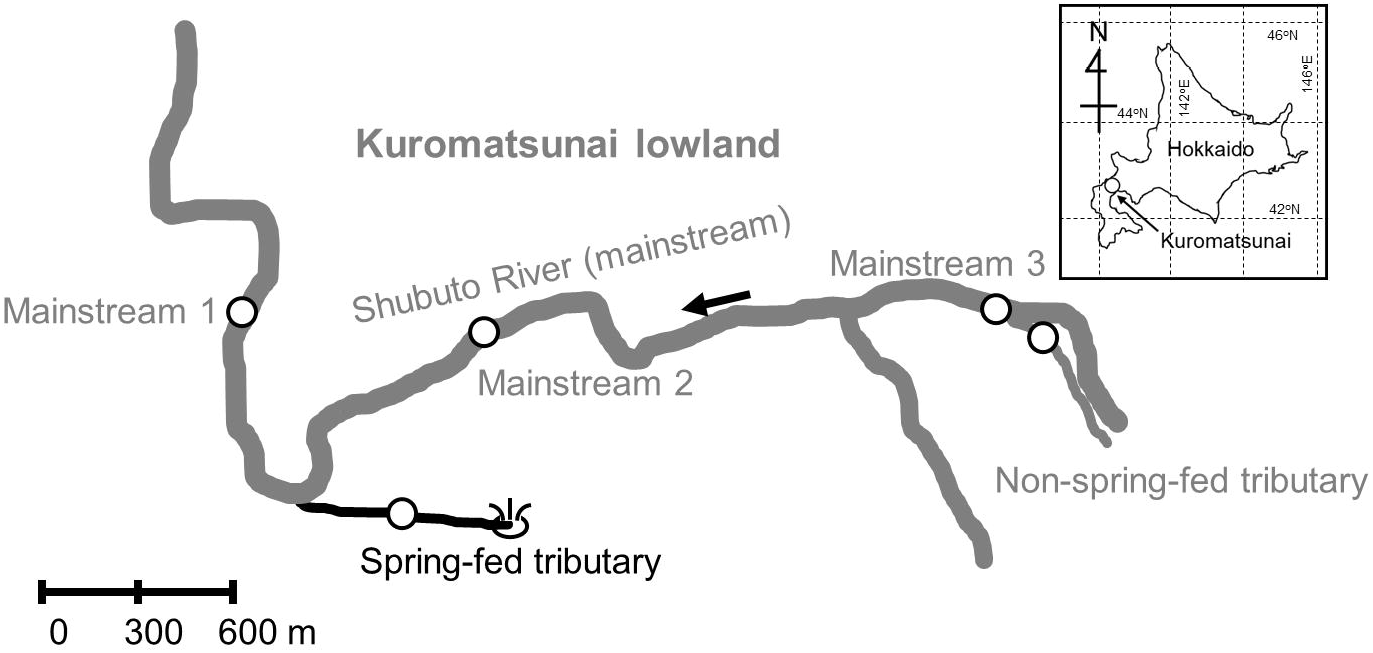
Study site location within the Shubuto River system in Kuromatsunai, Hokaido, Japan. Open circles indicate the study reaches.

**Fig. 2.**
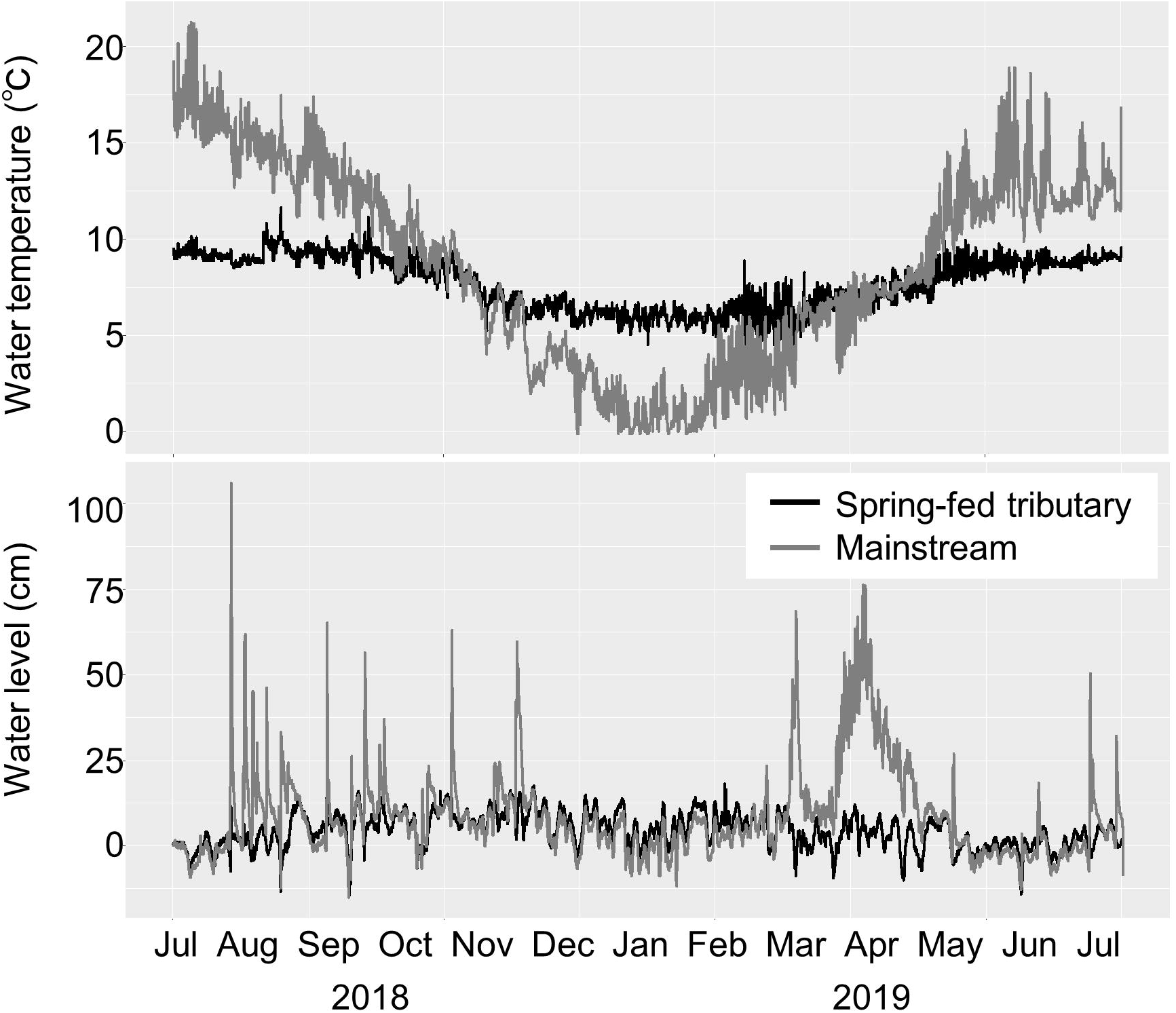
Flow and temperature regimes of the spring-fed tributary and adjacent mainstream. Water levels were initially set at 0 cm at the onset of the monitoring.

The lowland region of the Shubuto River basin includes areas of groundwater discharge such as flowing wells, seepages, and springs. The spring-fed tributary assessed in this study is fed by a significant amount of groundwater discharge (approximately 0.1 m^3^/s) that is directly connected to the Shubuto River (Sakai et al. 2020). This tributary has stable flow regime and water temperature compared to adjacent mainstream (Fig. 2). We further selected a non-spring-fed tributary and three mainstreams that also drained into the lowland area (Fig. 1). The spring-fed and non-spring-fed tributaries were near to each other and had similar flow directions. Three mainstream areas were identified below the confluence with the spring-fed tributary (mainstream 1), between the confluences with the spring-fed and non-spring-fed tributaries (mainstream 2), and below the confluence with the non-spring-fed tributary (mainstream3) (Fig. 1). We used 40-m sections (reaches) in each study site to assess macroinvertebrates and habitat characteristics. The mainstream reaches had a typical fluvial geomorphology of Japanese midstream with riffle-pool structures whereas the tributary reaches rarely had heterogeneous structures like riffle-pools, step-pools, or cascades.

Although the river system has the only one typical spring-fed stream in the lowland area, similarly to other river systems in clastic geology, assumable unique habitat and community structures should be important for assessing local biodiversity even if focal river systems have limited numbers of spring-fed streams. This limitation may have hindered conceiving robust study design and advances in studies on ecosystem functions of spring-fed streams draining outside carbonate geologies. This study can therefore fill an important knowledge gap to understand the function of sparsely distributed spring-fed streams in structuring local biodiversity.

### Macroinvertebrate assemblage

We used Surber nets (25 × 25 cm, 0.5-mm mesh) to sample macroinvertebrates from five randomly placed quadrats in each reach in June 2016. This period was chosen to avoid seasons of snow accumulation, snowmelt, and primary insect emergences. Each sample was collected along center lines of the streams to obtain representative assemblages. The sampling points for the mainstream reaches categorized as riffles, but those for the tributary reaches did not have any clear riffle nor pool structures. Samples containing macroinvertebrates, detritus, and inorganic sediments were placed in a tray and the macroinvertebrate and detritus portions were rinsed into unwoven bags, and immediately preserved in 70% ethanol. Macroinvertebrates were sorted and identified to the lowest possible taxonomic level using a stereomicroscope (SZ61, Olympus, Tokyo, Japan) and keys published by Kawai and Tanida (2018) and Merritt et al. (2008). Following identification, we classified macroinvertebrates by life-form type (burrowers, sprawlers, clingers, and swimmers) and functional feeding group (shredders, collector-gatherers, collector-filterers, predators, and grazers) based on information from Takemon (2005) and Merritt et al. (2008) as well as our own unpublished data.

### Habitat characteristics

Stream width was measured with a measuring tape under base flow conditions at five points in each study reach. We measured water depth and current velocity (repeated 5-s measurements) using a folding scale and an electromagnetic current meter (VE10, Kenek, Tokyo, Japan), respectively, at 15 points in each study reach during macroinvertebrate sampling. Hemispherical photographs were taken in June 2016 at 10 m intervals along the 40 m reaches (five points) to estimate canopy openness. Photographs were taken with a digital camera (D40, Nikon, Tokyo, Japan) equipped with a fisheye lens (4.5 mm F2.8 EX DC Circular Fisheye HSM, Sigma, Kanagawa, Japan) that was fixed horizontally at 1 m above the streambed. Canopy openness (%) was estimated using CanopOn2 software (Takenaka 2009) from these photographs.

We applied the methods of Sakai et al. (2013) to estimate fine sediment deposition in each study reach by visually estimating the percent cover of fine sediment (< 2 mm) on the streambed in 40×40 cm quadrats with 20 rectangles separated by wires (5% increments, ten quadrats per reach). We also estimated the periphyton biomass on upper exposed surfaces of five submerged large cobbles in each study reach. Periphyton was collected by brushing material from a 4 cm^2^ patch on the upper cobble surface and filtering the resulting slurry through pre-combusted glass-microfiber filters (Whatman GF/F, Maidstone, United Kingdom). Prior to periphyton sampling, each cobble was thoroughly rinsed in stream water to remove attached fractions of nonalgal materials as much as possible. Periphyton dry mass and ash-free dry mass (AFDM) were determined by drying the periphyton samples in an oven (FC-610, Advantec, Tokyo, Japan) at 70°C for 24 h and then weighing them, followed by combustion in a muffle furnace (FO300, Yamato Scientific Co., Ltd., Tokyo, Japan) at 500°C for 2 h and re-weighing. Detritus samples remained after macroinvertebrate collections and identifications (five quadrat samples per reach) were dried at ambient temperature for three days, followed by oven drying at 70°C for three days and combustion in a muffle furnace at 500°C for 4 h. The AFDM of periphyton and detritus samples were used as measures of periphyton biomass and standing stock of detritus (mg/cm^2^), respectively.

### Statistical analyses

To test for differences in environmental variables, total abundance and taxa richness of macroinvertebrates, and abundance of the most dominant taxa (> 1% of the individuals in all samples) among the three stream types (spring-fed and non-spring-fed tributaries and mainstream), we first used one-way analysis of variance (ANOVA) with permutation test (Legendre 2007) to test for a significant difference of the response in at least one stream type. When significant, we used the same permutation ANOVA test for *post hoc* comparisons among the pairs of stream types. The permutation test used allowed us to correctly handle the unbalanced design of the study (mainstream has three times more observations than the tributaries). We adjusted *p*-values for multiples tests using the false discovery rate correction (Benjamini and Hochberg 1995), to avoid inflated type I error rates.

To visualize differences in macroinvertebrate assemblage structures among stream types and associated environmental variables, we performed a principal coordinate analysis (PCoA) based on the Chao index (Chao et al. 2005), followed by a permutational multivariate analysis of variance (PERMANOVA), testing for significant differences in macroinvertebrate assemblage structure among the stream types. Chao index was calculated from the total numbers of individuals of each taxon in each quadrat. We then superimposed *a posteriori* the environmental vectors (fine sediment deposition, canopy openness, current velocity, stream width, water depth, standing stock of detritus and periphyton biomass) using the weighted averages of the species, and environmental variables (Borcard et al. 2018). The PERMANOVA was tested using 999 permutations. In order to test for the presence of significant patterns of spatial autocorrelation in the macroinvertebrate communities across our study reaches, we generated spatial eigenvectors using Moran’s eigenvector maps (MEM; Dray et al. 2006), a powerful multivariate approach allowing the detection of multiscale spatial structures in ecological data. To obtain the spatial eigenvectors, we used a spatial weighting matrix based on a Gabriel graph and weighted by a decreasing linear function of the distance among streams (details in Bauman et al. 2018a), following Bauman et al. (2018a, b). A redundancy analysis of the reach-scale community matrix was performed against the whole set of spatial eigenvectors associated to positive spatial autocorrelation patterns, for each spatial weighting matrix, and was tested by permutations (9,999).

We additionally tested whether some of the macroinvertebrate taxa were significant indicators of some of the stream types, that is, displaying both high fidelity and specificity to one of the habitats. To do so, the *IndVal* index was calculated for all species in the three stream types (Dufrêne and Legendre 1997) and was tested using 10,000 Monte Carlo permutations. All statistical analyses were performed in R version 3.6.3 (R Core Team 2020) using the packages *rcompanion* (Mangiafico 2020), *adespatial* (Dray et al. 2020), *vegan* (Oksanen et al. 2019), *indicspecies* (Cáceres et al. 2020) and the function *anova*.*1way* (Legendre 2007).

## Results

Stream width and water depth were significantly greater in mainstream reaches and smallest in the non-spring-fed tributary (Table 1). Correspondingly, canopy openness was relatively greater in the mainstreams compared to the tributaries. Current velocity was significantly slower in the spring-fed tributary and faster in the mainstreams (Table 1). The percent cover of fine sediment was significantly higher in the spring-fed tributary; mean cover was approximately 11.4 times greater than those of the mainstreams and 3.8 times greater than that of the non-spring-fed tributary (Table 1). The periphyton biomass was not significantly different among the reaches, whereas the standing stock of detritus was greatest in the spring-fed tributary (Table 1).

**Table 1.**
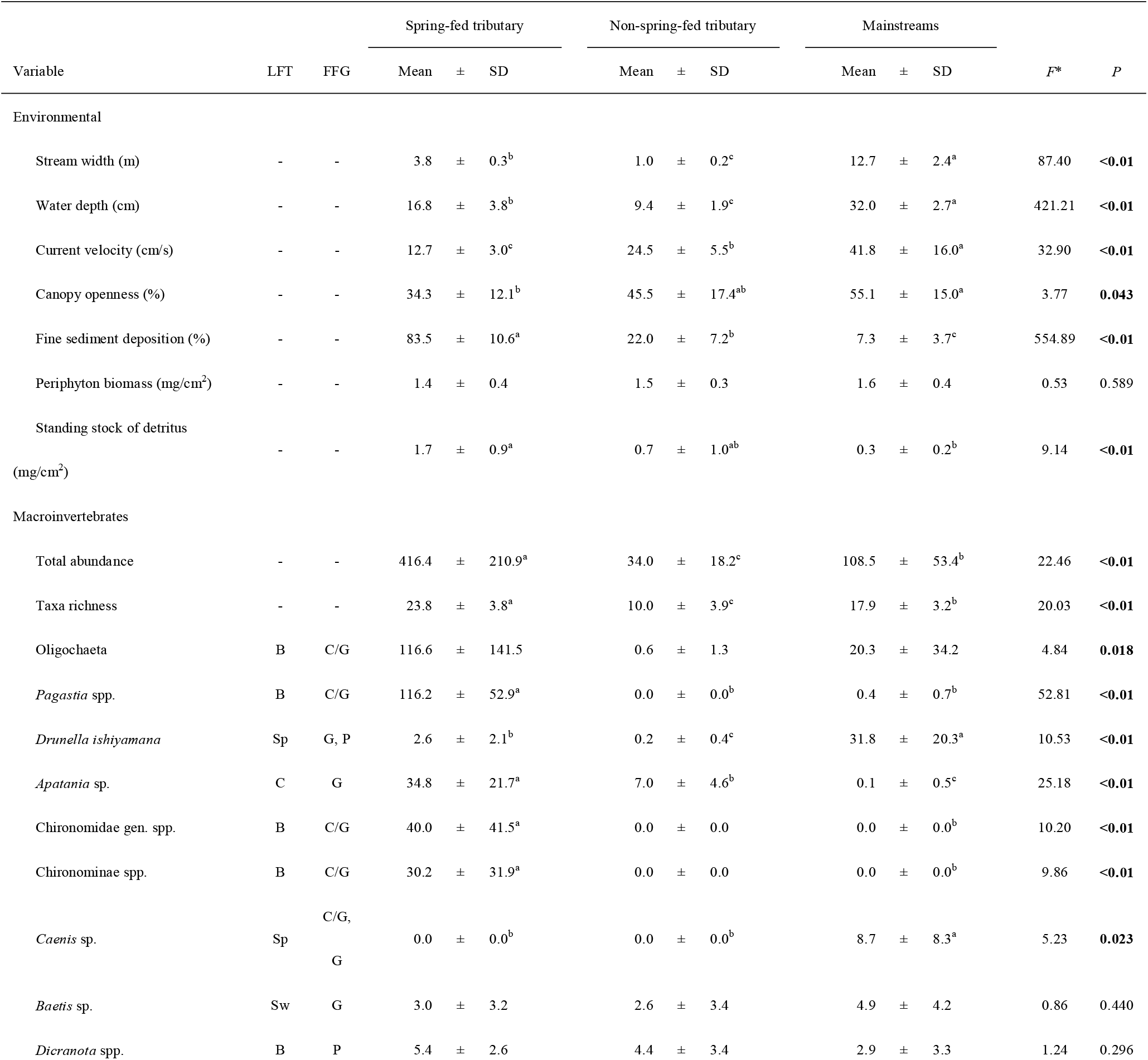
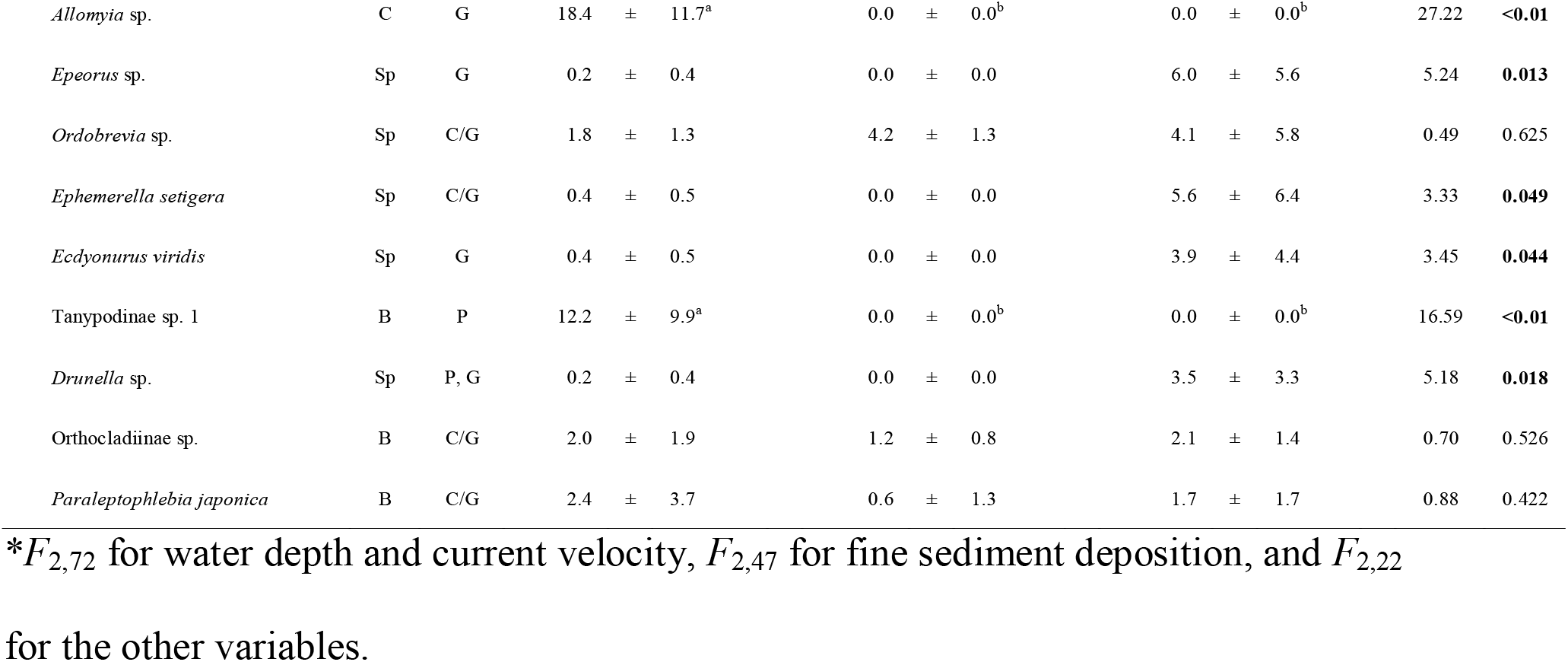
Environmental characteristics, total abundance, taxa richness, and the abundance of each dominant taxa in study reaches. LFT: life form type, B: burrower, Sp: sprawler, C: clinger, Sw: swimmer. FFG: functional feeding group, C/G: collector-gatherer, P: predator, G: grazer. Bold characters indicate statistically significant difference among stream types by one-way analyses of variances with permutation tests (*P* < 0.05), and different alphabetical letters indicate statistically significant differences by *post hoc* pairwise tests (*P* < 0.05). *P*-values are corrected for multiple tests (see Methods).

The total abundance and taxa richness of macroinvertebrates were both significantly greater in the spring-fed tributary and smaller in the non-spring tributary (Table 1; Supplementary Table S1). For life-form types, the four most abundant burrower taxa (Oligochaeta, *Pagastia* spp., Chironomidae gen. spp. and Chironominae spp.) and Tanypodinae sp. were more abundant in the spring-fed tributary. Sprawlers were typically the most abundant group in the mainstreams, except for *Ordobrevia* sp. (Table 1). For functional feeding groups, the four most dominant collector-gatherer taxa (Oligochaeta, *Pagastia* spp., Chironomidae gen. spp. and Chironominae spp.) were most abundant in the spring-fed tributary (Table 1). Excluding two trichopteran grazers that were dominant in the spring-fed tributary (*Apatania* sp. and *Allomyia* sp.), all grazer taxa reached their maximum abundance in the mainstreams. We noted that *Allomyia* sp. was only found in the spring-fed tributary (Table 1). In this study, some individuals of Chironomids could not be identified into subfamily or lower taxonomic levels, so that Chironomidae gen. spp. may potentially influence the total number of taxa, particularly in the spring-fed tributary that housed numerous Chironomid midges (Table 1).

The MEM analysis did not detect any significant patterns of spatial autocorrelation in the macroinvertebrate assemblage composition (*P* = 0.68), indicating spatial independence among the communities. The first two axes of the PCoA captured 97.6% of the total variation in the community Chao index matrix (Fig. 3), and the PERMANOVA indicated that macroinvertebrate assemblage structure significantly differed among the stream types (*P* = 0.001). After overlapping the environmental variables to the PCoA, the first PCoA axis indicated that the mainstreams presented greater water depth, stream width, periphyton biomass, current velocity and canopy openness than the spring-fed and non-spring-fed tributaries. The second PCoA axis mostly separated the non-spring-fed tributary from the spring-fed tributary, the two presenting very contrasted community composition as well as fine sediment deposition and standing stock of detritus (higher for the spring-fed tributary). The indicator taxa for the spring-fed tributary were mostly detritivores (shredders, collector-gatherers and collector-filterers) of Chironomidae, Nemouridae, Ceratopogonidae, Lepidostomatidae, Limnephilidae, Pisidiidae and Eusiridae, but also included one herbivorous trichopteran (*Allomyia* sp.) (Table 2). The indicator taxa for the mainstreams were mainly grazers (*Caenis* sp., *Epeorus* sp., *Drunella* sp., *Setodes* sp. and *Ecdyonurus viridis*) and partly contained predators (*Drunella* sp. and *Rhyacophila nipponica*), collector-gatherer (*Ephemerella setigera*) and collector-filterer (*Cheumatopsyche infascia*) whereas those for the non-spring-fed tributary were collector-gatherer and/or grazer (*Agapetus budoensis*) and collector-filterer (Simuliidae gen. spp.) (Table 2).

**Table 2.** Statistically significant indicator taxa for each stream type (*P* < 0.05; adjusted *P* for multiple tests). LFT: life form type, B: burrower, C: clinger, Sp: sprawler, Sw: swimmer. FFG: functional feeding group, C/G: collector-gatherer, G: grazer, P: predator, S: shredder, C/F: collector-filterer.

| Taxon | LFT | FFG | Stream type | *IndVal* | *P* |
| --- | --- | --- | --- | --- | --- |
| Chironomidae gen. spp. | B | C/G | Spring-fed tributary | 1.000 | 0.0001 |
| *Allomyia* sp. | C | G | Spring-fed tributary | 1.000 | 0.0001 |
| Tanypodinae sp. 1 | B | P | Spring-fed tributary | 1.000 | 0.0001 |
| *Pagastia* spp. | B | C/G | Spring-fed tributary | 0.998 | 0.0001 |
| *Nemoura* sp. | Sp | S | Spring-fed tributary | 0.995 | 0.0001 |
| Chironominae spp. | B | C/G | Spring-fed tributary | 0.894 | 0.0009 |
| Tanypodinae sp. 2 | B | P | Spring-fed tributary | 0.894 | 0.0009 |
| Ceratopogonidae gen. spp. | B | C/G | Spring-fed tributary | 0.894 | 0.0009 |
| *Lepidostoma japonicum* | Sp | S | Spring-fed tributary | 0.824 | 0.0052 |
| *Nothopsyche* sp. | Sp | C/G, S | Spring-fed tributary | 0.775 | 0.0101 |
| *Pisidium japonicum* | B | C/F | Spring-fed tributary | 0.775 | 0.0080 |
| *Stenomoera yezoensis* | Sw | C/G, S | Spring-fed tributary | 0.764 | 0.0068 |
| *Agapetus budoensis* | C | C/G,<br>G | Non-spring-fed tributary | 1.000 | 0.0002 |
| Simuliidae gen. spp. | C | C/F | Non-spring-fed tributary | 0.764 | 0.0124 |
| *Caenis* sp. | Sp | C/G,<br>G | Mainstream | 0.966 | 0.0001 |
| *Epeorus* sp. | Sp | G | Mainstream | 0.950 | 0.0004 |
| *Drunella* sp. | Sp | G, P | Mainstream | 0.940 | 0.0006 |
| *Ephemerella setigera* | Sp | C/G | Mainstream | 0.899 | 0.0028 |
| *Setodes* sp. | Sp | C/G,<br>G | Mainstream | 0.856 | 0.0044 |
| *Ecdyonurus viridis* | Sp | G | Mainstream | 0.852 | 0.0154 |
| *Cheumatopsyche infascia* | C | C/F | Mainstream | 0.775 | 0.0193 |
| *Rhyacophila nipponica* | Sp | P | Mainstream | 0.730 | 0.0464 |

**Fig. 3.**
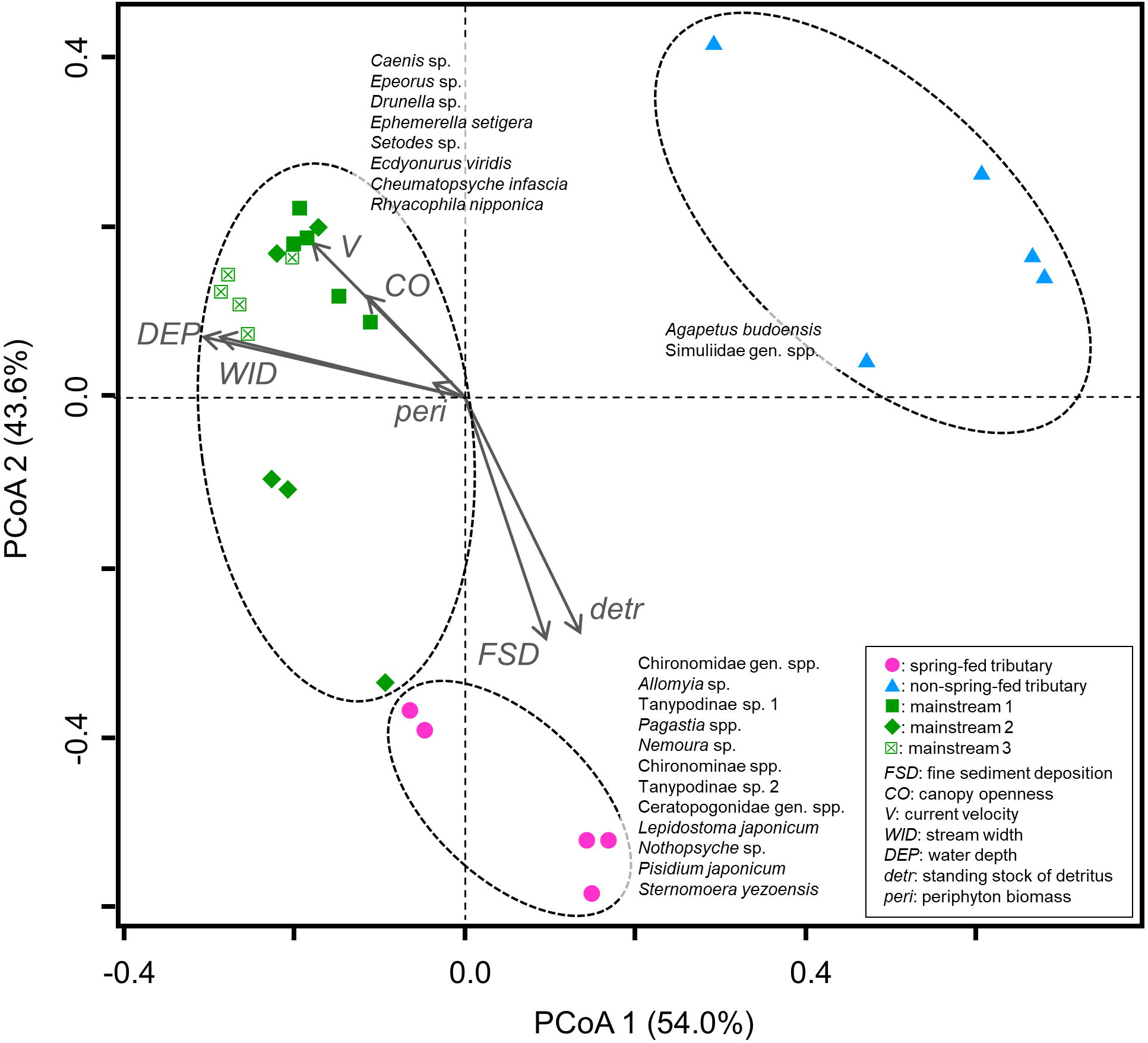
Results of the principal coordinate analysis based on the abundance of all macroinvertebrate taxa and the environmental variables. Significant indicator taxa for each stream type are shown nearby the plots.

## Discussion

Spring-fed streams in lowlands rarely have steep, deep channels (Sear et al. 1999) and their flow regimes are stable (Mattson et al. 1995; Lusardi et al. 2016). Sakai et al. (2020) showed that the spring-fed tributary assessed here maintained a stable flow regime even during rainfall events, implying that sediment runoff during floods likely occurs less in the tributary than in other non-spring-fed reaches. Because stable flow regime can settle fine particles in streambed (e.g., Kiffney et al. 2000), the significantly greater cover of fine sediment and higher standing stock of detritus observed in the spring-fed tributary are attributable to its stable flow regime. The observed slower current velocity in the spring-fed tributary could further contribute to the settlement of fine particles. The relatively closed canopy openness in the spring-fed tributary might additionally enhance accumulations of submerged detritus deriving from litter inputs.

Oligochaeta and most of the chironomid taxa were remarkably abundant in the spring-fed tributary. As a result, the total macroinvertebrate abundance was 3.8–12.2 times greater in this tributary than in the other reaches. The relative abundance of burrowers increases with fine sediment deposition (e.g. Zweig and Rabení 2001; Rabení et al. 2005), and our findings suggest that the fine substrate found in spring-fed streams provides attractive habitat for burrower species. Moreover, the stable flow regimes of spring-fed streams may relate to less disturbance to macroinvertebrates during flood events (e.g., Fritz and Dodds 2004; Robinson et al. 2004). Because the burrower species that dominated in the spring-fed tributary typically gather and feed on fine detritus (i.e., collector-gatherers), spring-fed environments clearly provide adequate food resources for this group. The assumable long retention of submerged detritus and associated consumptions and dominance of detritivores may thereby form detritus-based food webs in spring-fed streams. Meanwhile, the strong variation of macroinvertebrate abundances among the quadrats of a given stream type (e.g., Oligochaeta) suggest potential undetected microhabitat heterogeneity at finer spatial scale, within the relatively homogeneous habitats found in the spring-fed tributary.

In contrast to our results, previous studies have reported that spring-fed streams often have lush macrophytes (e.g., Sherwood et al. 2000; Katagiri et al. 2011) and are accordingly dominated by herbivorous invertebrates (Takemon 2010). Although obligate herbivores feeding on macrophytes are generally rare, macrophytes may supplement food resources for grazers that usually consume algal masses and provide spatially complex habitats. The streambed of the spring-fed stream we studied lacked macrophytes perhaps because of limited light conditions. Food web and habitat structures in spring-fed streams are probably site-specific and tightly associated with environmental factors that affect the growth of macrophytes.

Wider streams with more open canopies generally induce higher primary productivity (e.g., Vannote et al. 1980), but periphyton biomass was not significantly different between the mainstreams and tributaries in this study. However, ephemeropteran grazers significantly predominated in the mainstream reaches. This contradiction is attributable to greater occupancy by grazers in reaches with higher periphyton productivity. Feeding pressure of grazers is one of primary factors reducing periphyton biomass (Feminella et al. 1989; Hill et al. 1995), and reaches with higher productivity often have lower or similar periphyton biomass compared to those with lower productivity (Feminella and Hawkins 1995). Therefore, grazing-based food webs may be more pronounced in mainstreams and the dominance of grazer taxa and associated consumption of periphyton may be responsible for the lack of differences in periphyton biomass among reaches. The dominance of sprawlers in mainstreams is attributable to their adaptability to fast-current habitats.

In the PCoA, the mainstream 2 had somewhat similar macroinvertebrate assemblages to those in the spring-fed tributary. This result suggests that mainstream reaches may have more heterogeneous habitats compared to tributaries, and partially possess spring-fed-like macroinvertebrate communities even though the samples were collected from riffles. For example, patches with upwelling water within reaches can have such communities (Milner et al. 2001). Meanwhile, the macroinvertebrate assemblages in the non-spring-fed tributary were clearly different from those in the spring-fed tributary. Also, *Agapetus budoensis*, which was an indicator taxon of the non-spring-fed tributary, was only found there. This indicates that the non-spring-fed tributary also has unique habitats and macroinvertebrates assemblages compared to the spring-fed tributary and mainstreams. Although taxa richness was lowest in the non-spring-fed tributary, combinations of mainstream, spring-fed and non-spring-fed tributaries can be important for maintaining the beta and gamma diversities of the river network.

Given that the basal food resources and macroinvertebrate communities differed between the spring-fed tributary and the adjacent mainstream reaches, we suggest that the confluence of these streams may tangle the structure of the food web such that the abundant macroinvertebrates in the spring-fed tributary may subsidize predatory animals, such as fishes, inhabiting surrounding streams (cf. Uno and Power 2015). Moreover, the stable flow regime and water temperature in spring-fed streams provide refugia during flood events (Sakai et al. 2020) and thermal stresses (Inoue and Ishigaki 1968). Terui et al. (2018) reported that complex branches in river networks can increase metapopulation stability for fish. Spring-fed streams may also contribute to such stabilities in stream animal communities. Given that spring-fed habitats are ubiquitous and distributed across Japan, examining the ecosystem functions of these streams in nutrient cycling and as refugia is an important topic in river ecosystem conservation. Given that the climate crisis relates to extreme runoff events (Yin et al. 2018) and severe thermal stress (van Vliet et al. 2013), these streams may be vital to conservation efforts.

Two sprawler-grazer trichopteran species (*Allomyia* sp. and *Apatania* sp.) that can be considered crenobiont and/or crenophilous were dominant in the spring-fed tributary. In Japan, *Allomyia* species occur only in Hokkaido, and are preferentially found in cool spring-fed habitats (Nishimoto and Kuhara 2001; Kawai and Tanida 2018). *Allomyia* sp. was found only in the spring-fed tributary in this study, similarly to another study conducted in a montane spring-fed stream (Sueyoshi et al. 2014). At the global scale, many of the crenophilous invertebrates are stenothermal, which is related to their evolutionary origins (Sun et al. 2020), and the stable thermal environments of spring-fed streams may provide refugia for these coldwater taxa (Lusardi et al. 2016). The taxa that were dominant in the spring-fed tributary included stenothermal species, implying that unique assemblages may be found in such streams. This may be an important function of spring-fed streams; their unique habitats may contribute to the beta diversity of focal river networks (Reiss et al. 2016).

Intensive researches conducted in carbonate geologies revealed that locations of spring habitats are largely influential on local faunas. For example, faunal characteristics tend to be similar between alpine springs and headwaters (Wigger et al. 2015), and substrate and macroinvertebrate assemblage structures vary with altitudinal gradients (von Fumetti and Blatter 2017). Additionally, regions that receive small amount of precipitations need to consider flow permanence of lotic systems. Some studies evaluated faunal characteristics in between intermittent and perennial spring habitats indicated similarity between them in high flow seasons (Wood et al. 2005; White et al. 2018) but life cycle adaptations of some taxa in the intermittent habitats (Wood et al. 2005). In lowland regions, Reiss et al. (2016) reported that crenobiont taxa which can be considered as glacial relicts inhabit in spring habitats, corresponding to this study. Although intermittent spring habitats may be relatively less under the moist climate of Japan, a concern regarding impact of anthropogenic disturbance such as land modifications and climate change on spring habitats (Cantonati et al. 2012) is common across locations and geologies.

Our comparisons of spring-fed and non-spring-fed streams revealed unique habitat characteristics and macroinvertebrate assemblage structure in the spring-fed tributary. The macroinvertebrate assemblage was characterized by numerous burrowers and collector-gatherers, and possibly crenobiont and/or crenophilous taxa. This unique assemblage may therefore enhance the beta-diversity of this river network. Owing to the high cover of fine sediments, the abundance of burrower taxa was remarkably high in the spring-fed tributary. These taxa may further play significant roles as prey for carnivores in the surrounding area. Although spring-fed streams are sparsely distributed under clastic geology, ideal site selections that include multiple spring-fed and non-spring-fed streams would further improve our understandings how spring-fed habitats structure animal communities. Given that spring-fed streams may act as flow and temperature refugia (Sakai et al. 2020, Lusardi et al. 2016), further studies that examine availabilities of the refugia in spring-fed streams can inform biodiversity conservation in river ecosystems. Here, we highlight the ecosystem function of providing unique habitats and housing unique macroinvertebrate assemblages.

## Acknowledgments

Dr. Izumi Washitani provided invaluable comments on the study plan. We thank the field assistance by Dr. Kosei Takahashi, Mr. Hitoshi Saito, Nobuo Hatai and Kengo Ebihara.

## Availability of data and materials

The data that support the findings of this study are available from the corresponding author upon reasonable request.

## Authors’ contributions

MS, KI and DB conceived and designed the study. MS and KI conducted the field investigations, and MS and DB performed the statistical analyses. MS and KI wrote the first draft, and all authors contributed to the writing of the final manuscript.

## Ethical approval

Not applicable.

## Consent to participate

Not applicable.

## Consent for publication

Not applicable

## Code availability

Not applicable.

